# Myeloid A20 is critical for type-2 immune mediated helminth resistance

**DOI:** 10.1101/2023.09.05.556360

**Authors:** Ioanna Petta, Marie Thorp, Maarten Ciers, Gillian Blancke, Louis Boon, Tim Meese, Filip Van Nieuwenburgh, Andy Wullaert, Richard Grencis, Dirk Elewaut, Geert van Loo, Lars Vereecke

## Abstract

Protective immunity against intestinal helminths requires induction of robust Type-2 immunity orchestrated by various cellular and soluble effectors which promote goblet cell hyperplasia, mucus production, epithelial proliferation and smooth muscle contractions to expel worms and reestablish immune homeostasis. Conversely, defects in type-2 immunity result in ineffective helminth clearance, persistent infection and chronic inflammation. We identify A20 as an essential myeloid factor for the induction of type-2 immune responses against the intestinal parasite *Trichuris muris*. Myeloid cell-specific loss of A20 in mice (A20^myel-KO^) results in chronic *Trichuris muris* infection and intestinal inflammation. Myeloid A20 deficient mice are not able to induce anti-helmith type-2 immune responses while instead mount detrimental Th1/Th17 polarized immune responses. Antibody-mediated neutralization of the type-1 cytokines IFNγ, IL18 and IL12 prevents Th1/Th17 polarization and reestablishes Type-2 mediated protective immunity against *Trichuris muris* in A20^myel-KO^ mice. In contrast, the strong Th1/Th17 biased immunity in A20^myel-KO^ mice offers protection against *Salmonella* infection. We hereby identify A20 as an essential myeloid factor to initiate approriate adaptive immunity in response to infection, and to induce a balanced type-2 immune response against the intestinal parasite *Trichuris muris*.

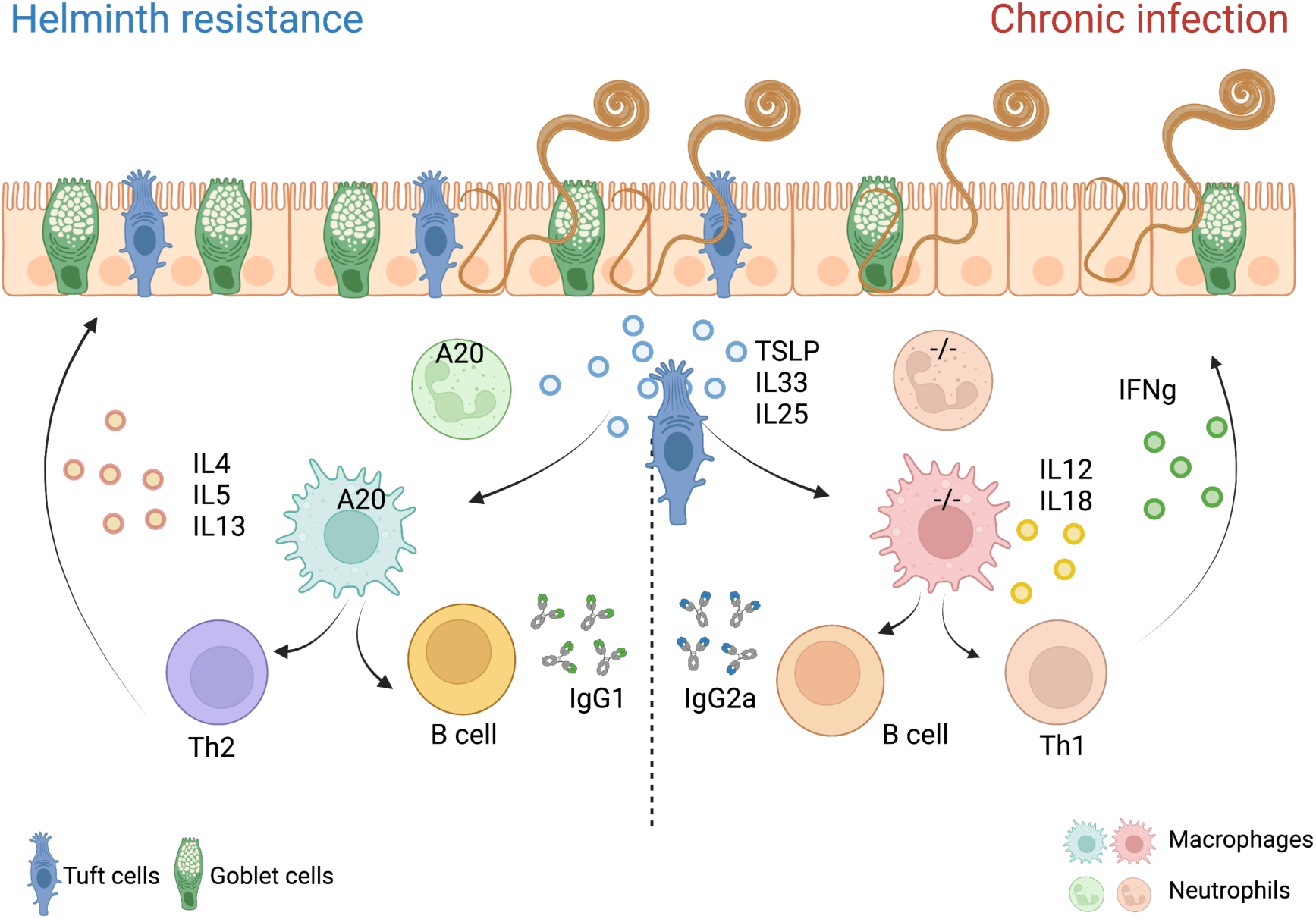

**Graphical abstract:** The clearance of gastrointestinal helmiths depends on type-2 immunity. Helminths interact with and damage intestinal tissue, which leads to the release of intracellular DAMPs and cytokines such as TSLP and IL33, and IL25 produced by epithelial cells. These factors may activate myeloid cells and ILC’s, which further activate T and B cells to mount effective Th2 responses and the secretion of IL4, IL5 and IL13 cytokines, as well as helminth-specific IgG1 immunoglobulins, leading to effective expulsion of the helminths. Deletion of A20 in the myeloid cells leads to enhanced secretion of type-1 cytokines, including IL12, IL18 and IFNγ, which impede type-2 immune-mediated helminth clearance and promotes chronic intestinal inflammation.

## Introduction

Infections with parasitic worms (helminths) affect more than a quarter of the world’s population, and are most prevalent in developing countries with poor hygiene and sanitation^1^. The soil-transmitted-helminths *Ascaris lumbricoides*, *Trichuris trichiura*, *Ancylostoma duodenale* and *Necator americanus* are responsible for most infections in humans and can cause severe morbidity^2^. In order to study immunity against helminth infections, most studies are done using the murine parasite *Trichuris muris*, which serves as a representative model for the human helminth *Trichuris trichiura,* which infects 465 million people worldwide^3^. Helminths are large parasites which cannot be removed by phagocytosis. Protective immunity against helminths involves complex host-parasite interactions and is characterized by a robust type-2 (Th2) immune response in an attempt to clear the helminths, reinforce the mucosal barrier and induce tissue repair. The helminth-induced immune response is initiated by intestinal damage and the release of specific cytokines and alarmins, including IL25, IL33 and TSLP, which can activate innate immune cells and prime the induction of adaptive Th2 immune responses^4^. Type-2 cytokines IL4, IL5 and IL13, produced by Th2 cells and innate immune cells including mast cells, eosinophils, neutrophils and type-2 innate lymphoid cells (ILC2s), induce effector mechanisms to expel the multicellular parasites, including IL13-mediated goblet cell hyperplasia and mucus production, IgG1 immunoglobulin production by B cells, smooth muscle contraction and epithelial proliferation. Macrophages play an important role in regulating anti-helminth immunity, as they trap and kill parasite larvae, but also contribute to tissue repair and the resolution of type-2 inflammation^5^. Upon helminth infection, macrophages adopt an ‘alternatively activated’ macrophage (AAM) phenotype, which involves the production of various effector molecules including Arginase-1 (Arg1) and Resistin-like protein alpha/beta (Relmα/β). Neutrophils also respond to helminths by producing factors which drive AAM-polarization, including IL13^6^.

A20 (also known as Tumor Necrosis Factor-induced Protein 3, TNFAIP3) is a negative regulator of various inflammatory signaling pathways induced by Pattern Recognition Receptors (PRRs, including TLR’s and NLR’s), cytokine receptors (IL11β, TNF) and T-cell receptors. The anti-inflammatory properties of A20 are commonly attributed to its ability to suppress inflammatory NF-κB signalling, but recent studies in mice have demonstrated that A20 primarily suppresses inflammation by preventing cell death^7^. A20 exerts its function by acting as a ubiquitin-editing and ubiquitin-binding protein, thereby interfering with downstream signaling events^8,9^. Myeloid-specific A20 deficiency in mice causes excessive inflammatory cytokine production and spontaneous arthritis development, which is characterized by hyperactive NLRP3 inflammasome activity and macrophage necroptosis^10,11^. Helminths or their products have been suggested to have therapeutic potential for treating autoinflammatory and autoimmune diseases including IBD and arthritis, by counterbalancing pathogenic immune pathways^12,13^. In this study, we identify A20 as an essential myeloid factor that orchestrates effective anti-helminth type-2 immunity and prevents chronic Th1/T17-mediated inflammation upon helminth infection.

## Results

### Myeloid A20 deficiency results in altered intestinal immune cell composition

Myeloid-specific A20 deficient mice (A20^myel-KO^) develop spontaneous arthritis with age and show signs of systemic inflammation^10^. However, they do not develop spontaneous intestinal inflammation (Sup. Fig. 2A-C and ^14^). To study intestinal immunity in more detail, we performed immune phenotyping of the colonic lamina propria immune cells in A20^myel-KO^ and wild-type (WT) littermate control mice using multi-panel flow cytometry, and show that myeloid-specific A20 deletion leads to the expansion of various myeloid immune cell subsets, including neutrophils, monocytes and macrophages, compared to WT littermate controls (Figure 1A,B). In addition, colonic lamina propria T cell subsets in A20^myel-KO^ mice are characterized by expansion of Th17 (CD4+RORγt+) cells. In contrast, A20^myel-KO^ mice have lower levels of colonic CD4+GATA3+ cells in steady state condition (Figure 1C,D). Similar changes are observed in the spleen, although with less pronounced changes in the T cell compartment (Sup. Fig. 1). Despite changes in leukocyte profiles in the colon of A20^myel-KO^ mice, histological analysis of colon tissue do not show clear signs of inflammation (Sup. Fig. 2B), as previously described^15^. Since A20^myel-KO^ mice develop spontaneous arthritis in the absence of intestinal inflammation, we compared the myeloid cell compartment in both colon and joints by performing bulk RNA sequencing on sorted myeloid cells (CD11b+) from colon lamina propria (Figure 1E-H, sup. Fig 2E) and joint (combined ankle and knee synovium) tissue (Figure 1I-L, sup. Fig. 2F) between A20^myel-KO^ and WT littermates in steady state conditions. The top-upregulated genes in colonic A20^myel-KO^ CD11b+ cells are involved in IFN signalling and in classical macrophage differentiation (M1), including *Cxcl9, Gbp5*^16^*, Saa3, Gzma* and *Nos2*^17^ Figure 1E, F). We also performed Gene Ontology (GO) pathway analysis, which revealed that expression profiles of colonic myeloid cells from A20^myel-KO^ mice are associated with macrophage tolerance, cytokine pathways, including IL12, IL23, IL22, IL17, IFNγ signalling, and TLR4 signalling (Figure 1G). Gene scoring analysis using an ‘M1 signature calculation’^18^ indeed showed a significantly higher M1 score in the A20^myel-KO^ colonic myeloid compartment compared to WT colonic myeloid cells (Figure 1H).

**Figure 1:**
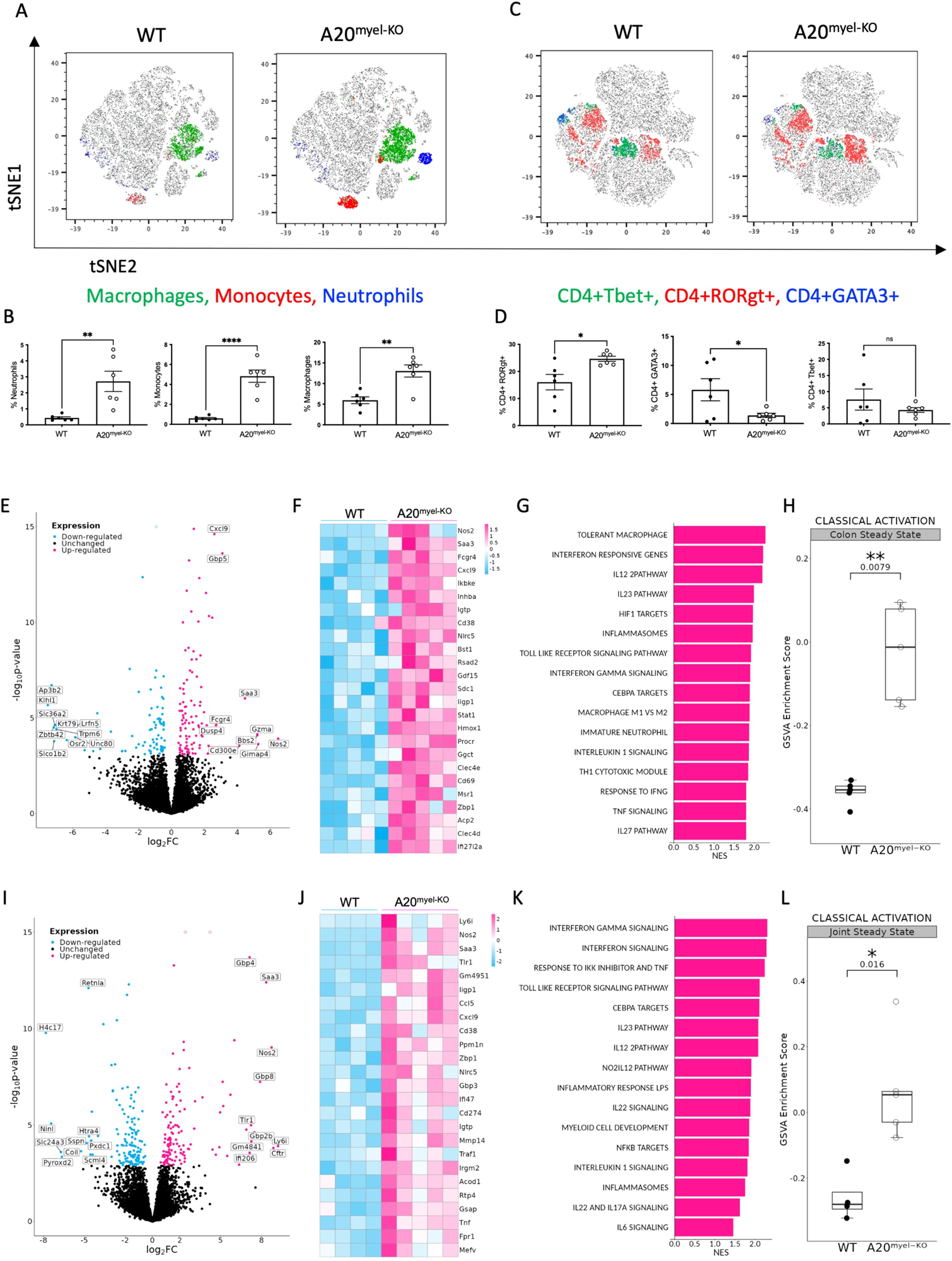
A20 deletion in myeloid cells affects immune cell composition and favors M1 macrophage polarization. (A, B) Flow cytometry analysis of colon lamina propria of WT (n=6) and A20^myel-KO^ (n=6) mice for the myeloid and (C, D) T cell composition, presented in tSNE plots (upper panels) and bar graphs (lower panels). All data are presented as means ± s.e.m. Statistical significance was determined by a two-sided Student’s t-test. E) Bulk RNA sequencing volcano plots presenting the differentially expressed genes in CD11b+ sorted cells of WT (n=5, blue) and A20^myel-KO^ (n=5, magenta) colon samples. F) Heatmap showing the top 25 M1 markers to be upregulated in A20^myel-KO^ colon compared to WT. G) Gene ontology pathway analysis showing the positively regulated pathways in A20^meyl-KO^ colon CD11b+ cells. H) Gene scoring analysis using an M1 signature consisting of 700 genes to compare expression in WT and A20^myel-KO^ colon. I) Bulk RNA sequencing volcano plots presenting the differentially expressed genes in CD11b+ sorted cells of WT (n=5, blue) and A20^myel-KO^ (n=5, magenta) joint synovium samples. J) Heatmap showing the top 25 M1 markers to be upregulated in A20^myel-KO^ synovium compared to WT. K) Gene ontology pathway analysis showing the positively regulated pathways in A20^meyl-KO^ synovium CD11b+ cells. L) Gene scoring analysis using an M1 signature to compare the expression in WT and A20^myel-KO^ synovium.

**Figure 2:**
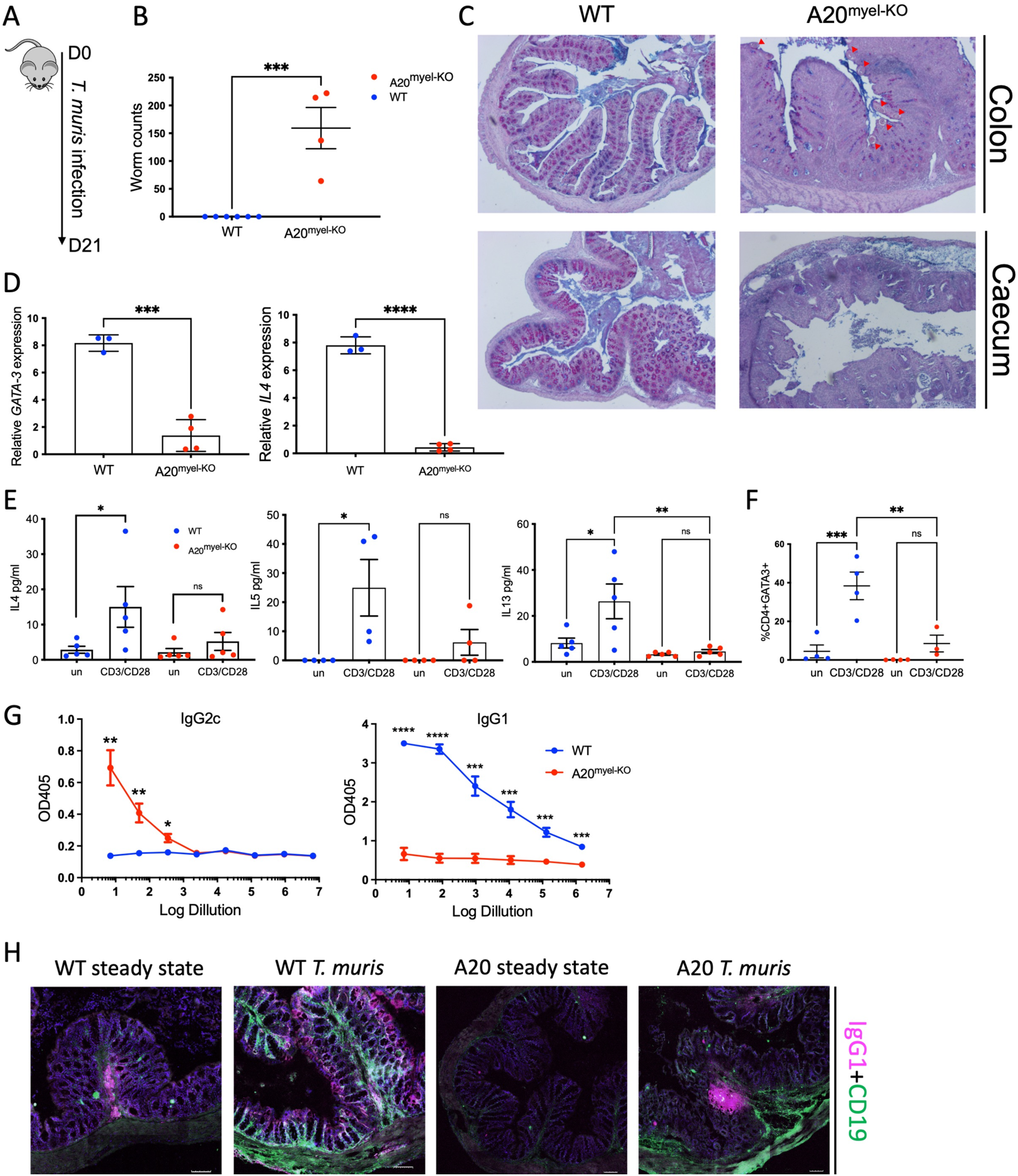
A20^myel-KO^ mice are susceptible to *Trichuris muris* infection. A) Infection scheme with Trichuris muris gastrointestinal helmith. B) Worm counts in the caecum of WT (n=6, blue) and A20^myel-KO^ (n=4, red) 21 days after infection with Trichuris muris. C) Representative images of AB/PAS staining of WT and A20^myel-KO^ colon (upper panel) and caecum (lower panel) 21 days after Trichuris muris infection. Scale bars 100μm. D) GATA3 and IL4 relative expression in WT (n=3) and A20^myel-KO^ (n=4) total colon lysates. E) Cytokine levels of IL4, IL5 and IL13 in the supernatant of restimulated with a-CD3/a-CD28 WT and A20^myel-KO^ mLN for 48h. F) CD4+GATA3+ levels determined by Flow cytometry analysis in restimuated mLN and of WT and A20^myel-KO^. G) ELISA for the measurement of IgG2c and IgG1 levels in the blood serum of WT and A20^myel-KO^ mice infected with Trichuris muris for 21 days. H) MACSima imaging for IgG1 (magenta) and CD19 (green) of WT and A20^myel-KO^ colon sections at steady state and upon infection with *Trichuris muris* for 21 days. Scale bars 100μm. All data are presented as means ± s.e.m. Statistical significance was determined by a two-sided Student’s t-test or by One-way anova for multiple comparisons.

RA patients show an enhanced M1/Th1 profile which is characterised by increased TLR signaling and secretion of inflammatory cytokines such as TNF, IL1β, IL18 and IFNγ^19^. Accordingly, analysis of A20^myel-KO^ synovial myeloid cells (CD11b+) revealed transcriptional upregelation of M1-genes, including different Gbp family members^20^, acute phase response markers (such as Saa3)^21^, TLR1 pathway genes^22^ and IFNγ-induced genes (such as *Gm4841*)^23^ (Figure 1I, J). Moreover, GO pathway analysis revealed that positively regulated pathways in the synovial myeloid compartment of A20^myel-KO^ mice include IFN-signaling, proinflammatory cytokine signaling (IL1, IL6, IL17, IL23), and NF-κB and inflammasome signaling (Figure 1K). Finally, we also identified a similar M1-gene signature scoring signaure for the joints, demonstrating that A20 deficient myeloid cells exhibit a prominent M1 phenotype compared to WT controls (Figure 1L). Together, these data indicate that A20 deletion in myeloid cells induces M1 priming in both intestinal and synovial immune cells, and is correlated with polarized Type1/3 adaptive immune polarization in steady-state.

### Myeloid A20 is essential for protective immunity against *Trichuris muris* infection

Given the reduced baseline Th2 cell numbers in colon and the macrophage polarization towards a pro-inflammatory M1 phenotype, we next evaluated the sensitivity of A20^myel-KO^ mice to an intestinal parasite infection, since helminth clearance critically depends on type-2 immune responses^24^. For this, adult A20^myel-KO^ and wild-type littermate control mice were inoculated with 200 embryonated *Trichuris muris* eggs by oral gavage and were sacrificed for analysis 21 days later (Figure 2A). While, wild-type mice induce robust type-2 immune responses and clear *T. muris* from the intestine to reestablish homeostasis within 21 days^25^, A20^myel-KO^ mice show defects in worm clearance and still contain around 150-200 *T. muris* parasites in their caecum 21 days post infection (Figure 2B). A20^myel-KO^ mice are chronically infected with *T. muris* and display chronic intestinal inflammation characterized by severe immune cell infiltration, tissue damage and edema (Figure 2C). We also show that GATA3 and IL4, two prototype type-2 immunity markers, are expressed in lower levels in A20^myel-KO^ colon lysates from *T.muris-*infected mice compared to WT littermates (Figure 2D). Restimulation of mesenteric lymph node (mLN) cells 21 days post infection (p.i.) with anti-CD3/CD28, induces production of type-2 cytokines, such as IL4, IL5 and IL13, in cells derived from wild-type mice but not in A20^myel-KO^ mLN cells (Figure 2E). WT-derived mLN are also characterized by higher levels of CD4+GATA3+ cells (Figure 2F). To further evaluate the induction of type-2 immunity in response to *T. muris* infection, we quantified *T. muris-*specific serum IgG1 (type-2 associated) and IgG2c (type-1 associated) immunoglobulins. We found IgG1 to be strongly increased in serum of wild-type mice after *T. muris* infection but not in serum from A20^myel-KO^ mice. In contrast, *T. muris-*specific type-1 associated IgG2c immunoglobulins are increased in A20^myel-KO^ mice while absent in wild-type mice (Figure 2G). We also show increased CD19 and IgG1 levels in WT colon sections after *T. muris* infection, which is not the case in A20^myel-KO^ colon tissue, in line with our findings for serum IgG1 (Figure 2H).

Immune profiling of colonic lamina propria immune cells after *T. muris* infection (21 days p.i.) confirmed the inability of A20^myel-KO^ mice to mount an effective Th2 response, in contrast to WT littermates. We detected increased immune (CD45+) cell infiltration in A20^myel-KO^colonic lamina propria, with increased levels of neutrophils, monocytes and macrophages (Figure 3A, B). In contrast, WT mice show increased levels of eosinophils, which are associated with Th2 immune responses (Figure 3A, B). In the T-cell compartment, WT mice show elevated levels of CD4+GATA3+ (Th2) cells, compared to A20^myel-KO^ mice. In contrast, A20^myel-KO^ mice show a skewed Th1/Th17 response, as CD4+Tbet+ and CD4+RORγt+ cells are shown to be increased (Figure 3C,D). Similar findings are observed in the spleen, confirming a systemic immune response (Sup. Fig. 4A,B). Interestingly, the phenotype of the recruited T regulatory cells (Tregs) is indicative of the general adaptive response, and we observe WT-derived Tregs to express higher levels of GATA3, while A20^myel-KO^-derived Tregs express higher levels of Tbet, supporting Th2 and Th1 polarisation, respectively (Suppl. Fig. 4D, 5). As previously described, Tbet expression in T regulatory cells supports their homesostasis and activity in a type 1-mediated inflammatory context^26^. Multiplex imaging on colon sections confirmed the increased number of colonic CD45 cells in WT tissue after infection with *T.muris*, however, immune cell infiltration is even more increased in colons of infected A20^myel-KO^ mice. Similar recruitment patterns are observed for CD11b+ myeloid cells and for CD64+ macrophages (Figure 3E). Innate Lymphoid Cells 2 (ILC2) have a key role in orchestrating anti-parasitic type 2 responses^27^. Imaging on colon tissue of *T. muris*-infected mice revealed lower expression of the ILC2 marker CD90.2, but increased expression of the ILC1 marker Nkp46^28^ in A20^myel-KO^ mice compared to WT littermates (Sup. Fig 6).

**Figure 3:**
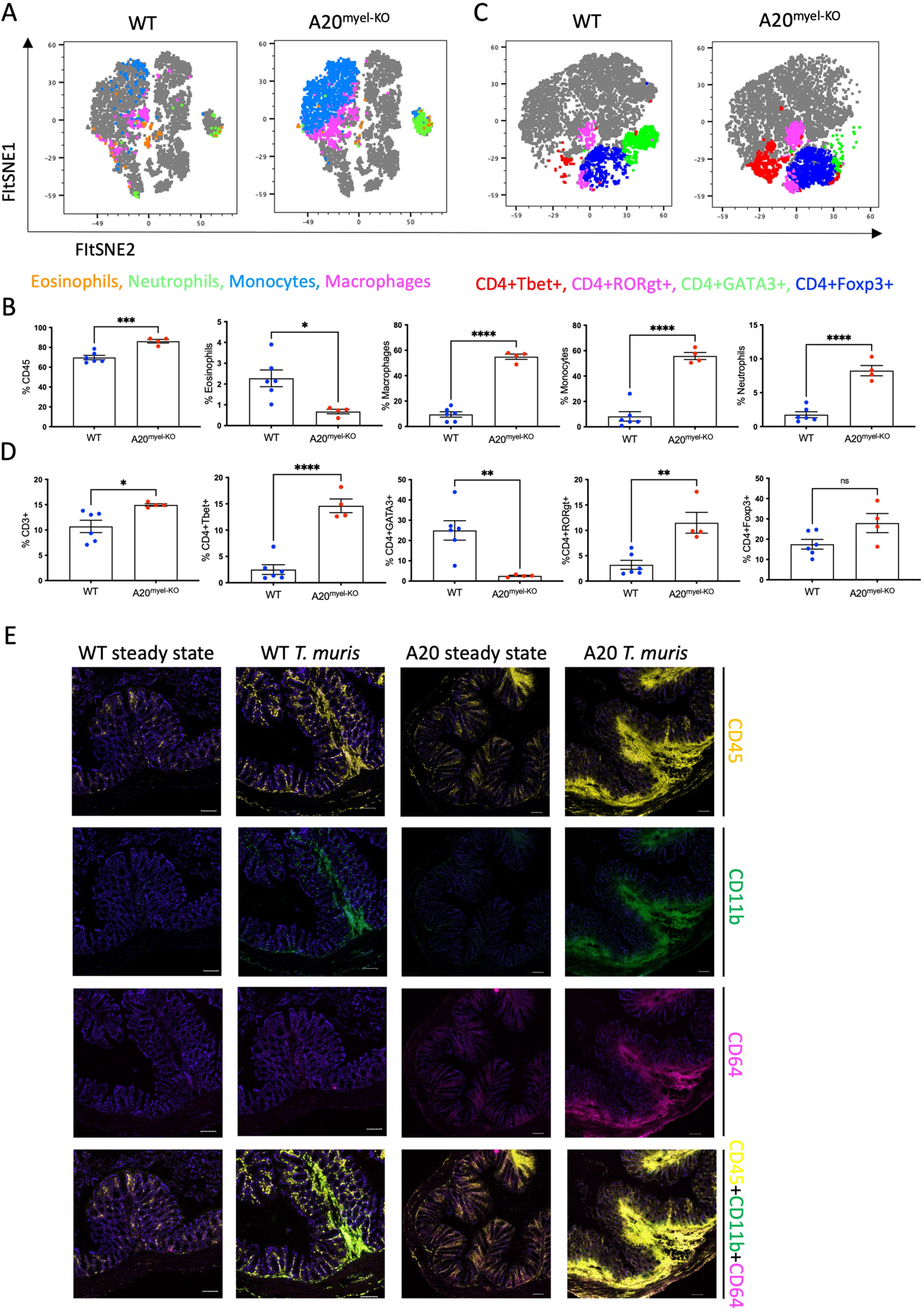
Deletion of A20 in myeloid cells hampers Th2 immunity and promotes Th1 response upon *Trichuris muris* infection. (A, B) Flow cytometry analysis of colon lamina propria of WT (n=6) and A20^myel-KO^ (n=4) mice infected with *Trichuris muris*, for the myeloid and (C, D) T cell composition, presented in tSNE plots (upper panels) and bar graphs (lower panels). E) MACSima fluorescent imaging for CD45 (yellow), CD11b (green) and CD64 (magenta) of WT and A20^myel-KO^ colon sections at steady state and upon infection with Trichuris muris for 21 days. Scale bar 100μm.

Finally, we performed RNA sequencing on sorted CD11b+ cells from colon of WT and A20^myel-KO^ mice after *T. muris* infection. Samples were isolated and processed 10 days after infection (D10), in order to investigate myeloid cell responses during the peak of infection and anti-helmith immune response. Interestingly, we observe a totally different expression profile for the sorted CD11b+ cells of both genotypes (sup. Fig 4C). In the A20 knockout myeloid cells we again observe the upregulation of markers related to classical macrophage activation such as *Nos2, CD300e*^29^*, Bmx*^30^ and *Plekhg4* (Figure 4A). GO analysis confirms the activation of pathways involved in IFN and TLR4 signaling, IL1β, IL12, IL22 sinaling, which are all related to macrophage polarization towards classical activation (Figure 4B). In contrast, in WT mice we observe the upregulation of markers related to alternative macrophage differentiation such as *CD163*^31^*, Nmb*^32^*, Cpeb1*^33^. GO analysis supports this notion, and indicates up-regulation of pathways related to IRF4 targets, PPARγ activation, IL4 and GATA3 signaling (Figure 4A, B). In order to further evaluate the difference in macrophage polarization, we compared the gene expression profiles of WT and A20^myel-KO^ mice to existing gene signature datasets of macrophage gene expression^18,34^. This confirms that WT mice show enhanced expression of genes involved in alternative macrophage activation, whereas A20^myel-KO^ mice show a clear polarization towards classical macrophage activation upon *Trichuris muris* infection (Figure 4 C,D). Moroever, Gene Set Enrichment Analysis (GSEA) confirmed the induction of an M1 phenotype, and the downregulation of an M2 phenotype in A20^myel-KO^ mice compared to WT mice (Figure 4E). Collectively, these data clearly indicate that upon *T.muris* infection, wild-type myeloid cells shift to an alternatively activated state, supporting proper type-2 immune polarization, pathogen clearance and resolution of inflammation. In contast, A20 deficient myeloid cells are highly polarized towards classical activation in steady-state, and this immune polarization is not altered upon *T.muris* infection. In conclusion, loss of A20 prevents myeloid cell polarization towards an alternatively activated state and subsequenctly impedes proper type-2 immune activation, which prevents parasite clearance and leads to chronic infection and inflammation upon helminth infection.

**Figure 4:**
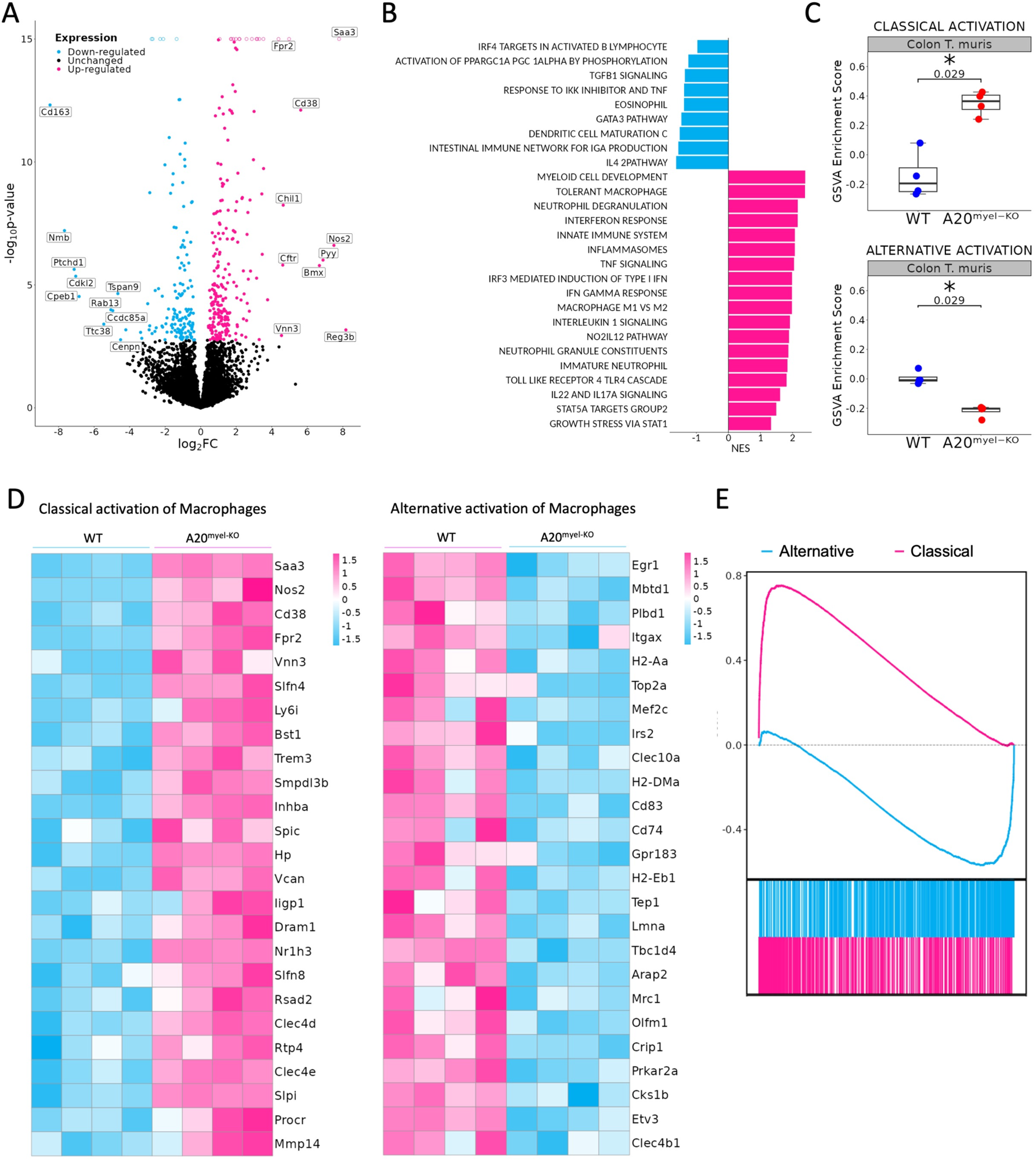
A20^myel-KO^ are characterized by M1 macrophage signature upon Trichuris muris infection. A) Bulk RNA sequencing volcano plots presenting the differentially expressed genes in CD11b+ sorted cells of WT (blue) and A20^myel-KO^ (magenta) colon samples of mice infected with *Trichuris muris* for 10 days. B) Gene ontology pathway analysis showing the positively regulated pathways in WT (blue) and A20^meyl-KO^ (magenta) colon CD11b+ cells of mice infected with Trichuris muris for 10 days. C) Gene scoring analysis using M1 and M2 signatures, consisting of 710 and 806 genes respectively to compare the expression in WT and A20^myel-KO^ colon upon Trichuris muris infection. D) Heatmap showing the upregulation of the top 25 markers related to classical activation of macrophages (M1) in A20^myel-KO^ mice compared to WT (left). Heatmap showing the upregulation of the top 25 markers related alternative activation of macrophages (M2) in WT compared A20^myel-KO^ mice (right). E) Gene set enrichment analysis (GSEA) plots visualizing the enrichment of classical (M1, upper panel) and alternative (M2, lower panel) activation of macrophages in A20^myel-KO^ and WT, respectively. Each plot features the running enrichment score (y axis) and shows the placement of the ranked differentially expressed genes for the respective transcriptomic signature (x axis).

### Type-1 cytokine inhibition promotes *Trichuris muris* resistance in A20^myel-KO^ mice

The Th1-associated cytokine IFNγ is a very potent antagonist of type-2 polarization, and is induced by upstream myeloid derived cytokines, including IL12 and IL18, which are elevated in A20^myel-KO^ mice (Sup. Fig. 2D). *Ex vivo* differentiated bone marrow-derived macrophages (BMDMs) of WT and A20^myel-KO^ mice were stimulated with *T. muris* antigens and analyzed at different time points. A20^myel-KO^ BMDMs express significantly higher levels of Th1 polarizing cytokines IL12 and IFNγ (Figure 5A, B) and secrete increased levels of IL18 (Figure 5C) in the supernatant upon *T. muris* antigen stimulation. In order to evaluate the role of IL12, IL18 and IFNγ in preventing an effective type-2 immune response upon *T. muris* infection in A20^myel-KO^ mice, we assessed the outcome of pharmacological inhibition of these cytokines *in vivo*. To this end, A20^myel-KO^ mice were infected with *T. muris* and i.p. injected with neutralizing antibodies against IFNγ, IL12 or IL18 versus isotype control antibody, every 48h for a total of 21 days (Figure 5D,E). Interestingly, depletion of either one of these three cytokines leads to a significant reduction in the worm counts in the caecum compared to isotype-antibody-treated A20^myel-KO^ mice, with most potent reduction in anti-IFNγ-treated mice (Figure 5E). We further determined the levels of IgG1 and IgG2c antibodies, as immunoglobulin markers of Th2 and Th1 responses, respectively. We observed that anti-IFNγ and anti-IL12 treatment had the most prominent effect in inducing a Th2-associated IgG1 antibody response, as efficient as in WT mice, although all A20^myel-KO^ mice retain increased levels of IgG2c (Figure 5F,G). Histological analysis confirms that inflammation is reduced in the colon of A20^myel-KO^ antibody-treated groups (Figure 5H). In conclusion, we have shown that A20^myel-KO^ macrophages overproduce type-1 inducing cytokines upon recognition of *T. muris* antigens, and that neutralization of these cytokines can re-establish effective anti-helminth type-2 immunity in A20^myel-KO^ mice.

**Figure 5:**
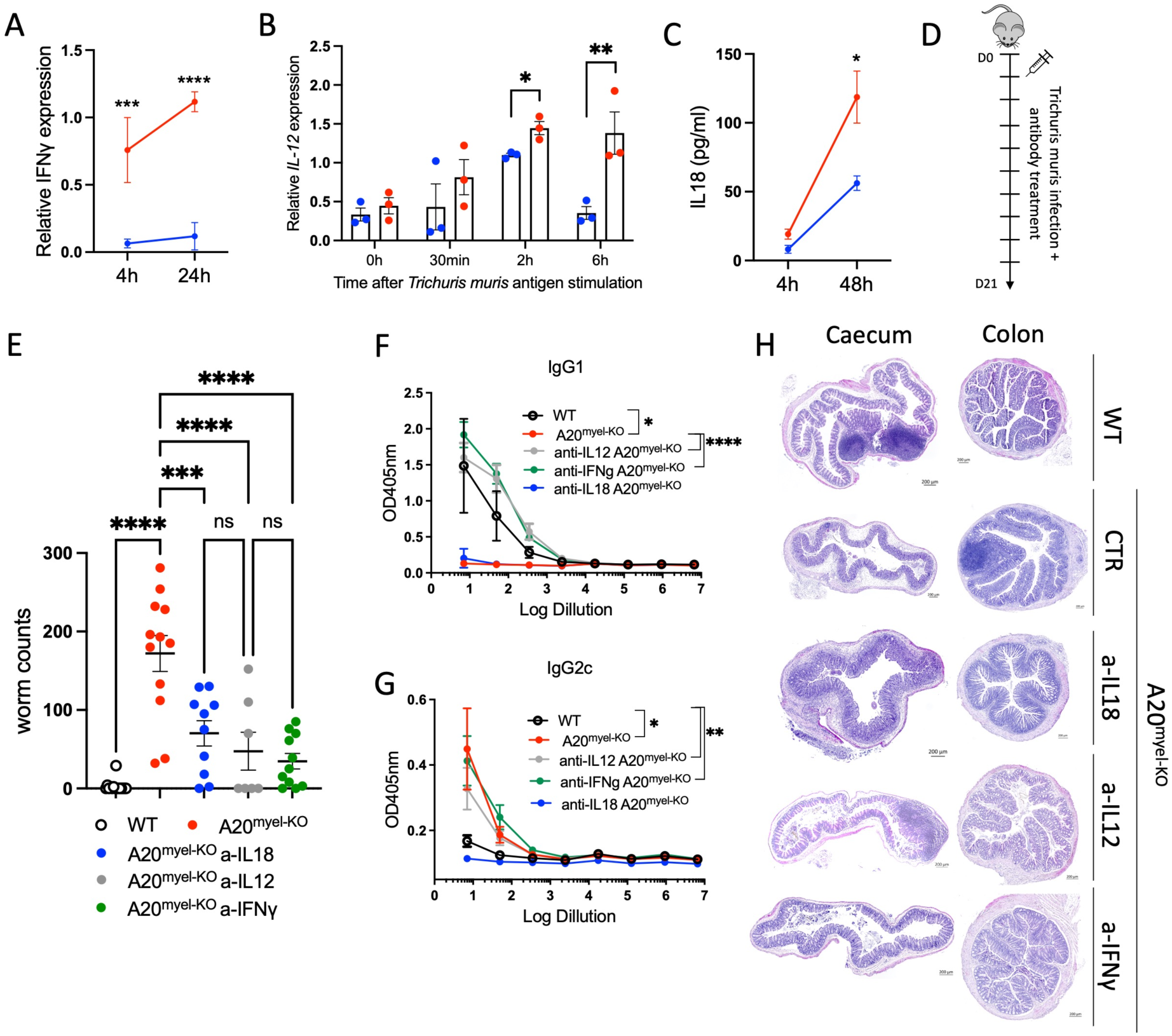
Blocking Th1-related cytokines reverse susceptibility of A20^myel-KO^ to *Trichuris muris*. A) Relative IFNγ expression in BMDMs derived from WT (blue) and A20^myel-KO^ (red) stimulated with *Trichuris* antigen for 4h and 24h. B) Relative expression of IL12 in WT (blue) and A20^myel-KO^ (red) derived BMDMs stimulated for 0min, 30min, 2h and 6h with *Trichuris* antigen. C) IL18 levels in the supernatant of WT (blue) and A20^myel-KO^ (red) derived BMDMs, stimulated with *Trichuris* antigen for 4h and 48h. D) Infection and treatment scheme with Trichuris muris and neutrilising antibodies against IFNγ, IL12 and IL18. E) Worm counts in the caecum of WT (n=15, open dots), A20^myel-KO^ (n=12, red), A20^myel-KO^+a-IL18 (n=10, blue), A20^myel-KO^+a-IL12 (n=7, grey) and A20^myel-KO^+a-IFNγ (n=11, green) 21 days after infection with *Trichuris muris*. The graph represents combined data of 3 independent experiments. F) ELISA for IgG1 and G) IgG2c levels in the blood serum of WT and A20^myel-KO^ mice treated as in B. Significance refers to log dilution 1. Η) Representative images of AB/PAS staining of caecum and colon sections of WT and A20^myel-KO^ mice infected with Trichuris muris and treated with neutralizing antibodies, as in B. Scale bars 200μm. Data are presented as means ± s.e.m. Statistical significance was determined by a two-sided Student’s t-test or by One-way anova for multiple comparisons.

### A20^myel-KO^ mice have improved resistance to Salmonella typhimurium infection

Given the strong Th1 immune profile of A20^myel-KO^ mice, we finally evaluated the sensitivity of A20^myel-KO^ mice and WT littermate controls to infection with *Salmonella typhimurium,* as a model for type-1 dependent bacterial infection. WT and A20^myel-KO^ mice were infected by oral gavage with 10^7^ CFU/mouse *S. typhimurium* for 8 days, and disease progression was assessed. A20^myel-KO^ mice show increased resistance against *S. typhimurium* infection compared to WT littermates, as they show increased survival (Figure 6A). Moreover, A20^myel-KO^ mice are able to restrict bacterial systemic dissemination and inflammation, as confirmed by significantly lower numbers of viable *S. typhimurium* derived from liver and spleen tissue (Figure 6B) and by significantly decreased levels of TNF in serum (Figure 6C). Histological analysis of spleen tissue of WT and A20^myel-KO^ mice showed a decrease of the lymphoid tissue (white pulp) over red pulp in WT compared to A20^myel-KO^ spleens (Figure 6D). Moreover, the liver of infected WT mice shows increased immune cell infiltration and necrotic areas compared to the liver of A20^myel-KO^ (Figure 6E). Although we do not observe differences in body weight and temperature between the two genotypes (Sup. Fig. 7C), A20^myel-KO^ are able to clear the *Salmonella* infection more effectively compared to WT littermates, which can be attributed to the enhanced type-1 immunity of A20^myel-KO^ mice.

**Figure 6:**
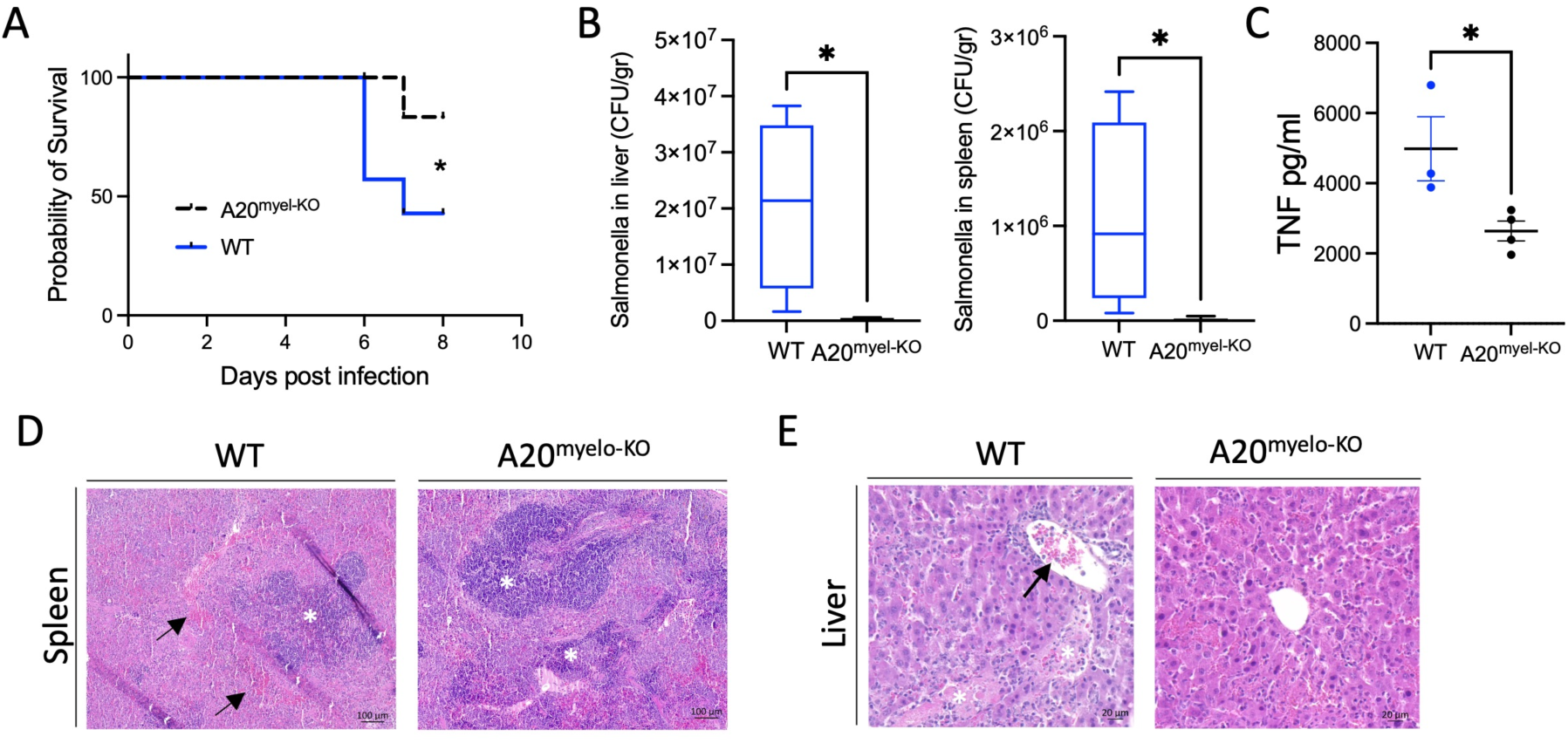
A20^myel-KO^ mice are protected against Salmonella Tiphimurium infection. A) Survival curve of WT (n=7) and A20^myel-KO^ (n=6) mice during 8 days of infection with *Salmonella Typhimurium*. Staistical analysis is performed by Gehan-Breslow-Wilcoxon test. B) CFU counts of retrieved Salmonella Tiphimurium from liver and spleen of infected WT (n=4) and A20^myel-KO^ (n=5) 8 days post infection. C) ELISA for TNF levels in the blood serum of WT (n=4) and A20^myel-KO^ (n=5) 8 days post infection. Data are presented as means ± s.e.m. Statistical significance was determined by a two-sided Student’s t-test. D) Representative images of H&E staining of spleens of infected WT and A20^myel-KO^ mice. Black arrows indicate hepatopoiesis (red pulp) and white asteric indicates the lymphoid tissue (white pulp). Scale bar 100μm. E) Representative images of H&E staining of liver of infected WT and A20^myel-KO^ mice. The black arrow indicates immune cell infiltration and white asterics indicate the necrotic areas. Scale bar 20μm.

### Discussion

A20 is a potent anti-inflammatory protein and its role in immune regulation has been extensively documented^35^. In this study, we report the role of A20 in macrophage polarization and show for the first time that its expression in myeloid cells is indispensable for the induction of effective Th2 adaptive immunity in response toinfection with the gastrointestinal helminth *Trichuris muris*.

The role of A20 in myeloid and T cells cells has been studied in an array of disease contexts. Deletion of A20 in myeloid cells in mice leads to spontaneous development of an erosive polyarthritis resembling rheumatoid arthritis in humans^11^. Follow-up studies have shown that the arthritic phenotype of A20^myel-KO^ mice is caused by increased Nlrp3 inflammasome activation, leading to increased secretion of IL1β and IL18^10^. Moreover, A20 was shown to prevent inflammasome-dependent arthritis by inhibiting macrophage necroptosis^36^. A20^myel-KO^ were also shown to be resistant to diet-induced obesity and insulin resistance due to the expansion of CD206+ adipose tissue macrophages (ATMs), although these ATMs have a gene signature indicative of classical pro-inflammatory macrophage activation^37^.

We showed that A20 deletion in myeloid cells prediposes intestinal myeloid cells towards a pro-inflammatory phenotype and M1 polarization in steady-state conditions, and drives subsequent Th1 immune responses in response to *Trichuris muris* infection. This in contrast to the WT control mice that efficiently induce Th2 immunity and clear the infection. Of note, *Gzma,* which we demonstrate to be upregulated in A20^myel-KO^ colon CD11b+ cells at steady state, was previously shown to be associated with chronic helminth infection to *Litosomoides sigmodontis,* and Granzyme A deletion *in vivo* was shown to promote Type-2 immunity^38^. Recently, A20 was also shown to control type-2 immunity in the lung. Deletion of A20 in macrophages was reported to suppress IL33–dependent expansion of lung ILC2s, type 2 cytokine production, and eosinophilia ^39^. These data are in line with our findings supporting an impaired intestinal type-2 immune response upon A20 deletion in colonic macrophages. Also, A20^myel-KO^ mice were shown to be protected against a lethal infection with influenza A virus^40^. Accordingly, we report here the increased resistance of A20^myel-KO^ mice against *Salmonella Typhimurium* infection, which also depends on pronounced type-1 immune responses.

In conclusion, we show that A20 deficiency not only leads to systemic inflammation and arthritis development, but also severely affects the sensitivity of mice to infection. While bacterial and viral infections, which depend on type-1 immunity, are cleared more effectively in the absence of A20, infections with parasites such as helminths, which depend on type-2 immunity, are compromised. These findings are important for patients with genetic defects in *TNFAIP3* (or functionally related genes). These patients show increased susceptibility to various inflammatory and autoimmune pathologies^41^, but may also have reduced resistance against parasite infections.

## Methods

### Animals

A20^myel-KO^ mice with a myeloid-cell-specific deletion of the A20/Tnfaip3 gene were previously described^11^. Mice were housed in individually ventilated cages at the specific pathogen-free animal facility of the Medical Research Building II at Ghent University Hospital. For our experiments, both males and females were used, evenly distributed over the experimental groups, as we observe no difference in resonse to *T. muris* and *S. Typhimurium* between both sexes. Ethics statement: All mice experiments were conducted according to the national Belgian Law (14/08/1986 and 22/12/2003, Belgian Royal Decree 06/04/2010) and European (EU Directives 2010/63/EU, 86/609/EEG) animal regulations. The described procedures were approved by the ethics committee of Ghent University Hospital (permit number LA1400410, approval ID 18-100).

### Propagation of Trichuris muris eggs

6-8 week old RaG2^-/-^ mice were infected with 200 *Trichuris* eggs by oral gavage. After 32-35 days the mice were sactificed and the caecum and proximal colon were isolated. The caecum and colon are cut open to expose the lumen and the worms placed in a 10cm petri dish containing PBS and with the help of forceps shake gently to remove feaces. The caecum was placed into another petri dish containing Dulbecco’s Modified Eagle Medium (DMEM) with 500 U/ml penicillin and 500 μg/ml streptomycin (P/S) and by using smooth curved forceps worms gently pulled out and placed in a well of a 6 well plate containing 5 ml of DMEM + P/S. This process was repeated for all of the infected mice. The plate was placed in a plastic container with a damp paper towel to maintain humidity and incubated at 37°C for 4 hr. After 4 hr worms were removed using smooth curved forceps and split into 2 new wells of the 6 well plate containing 5ml DMEM + P/S. The content of the original well was harvested and placed in a 15 ml falcon tube,spun tube at 3000 rpm and the supernatant removed and reserved as 4hr *T. muris* excretory and secretory antigen. This supernatant was used for mLN restimulation and BMDM treatment. The egg-containing pellet was resuspended in autoclaved tap water. The 6 well plate was placed back at 37°C overnight. Therafter, the worms were removed and the content of the wells harvested into a falcon tube and spun at 3000 rpm for 5 min. Thesupernatant was collected and reserved as 24h *T. muris* excretory and secretory antigen. This supernatant was used for *Trichuris* Ag specific antibody isotype ELISAs. The remaining egg-containing pellet was resuspended in autoclaved tap water and eggs embryonated at room temperature prior to storage at 4°C for future use.

### Infection with Trichuris muris helminth

Infection with Trichuris muris was performed by oral gavage of 200 embryonated *T. muris* eggs/mouse to 12 week-old WT and A20^myel-KO^ mice. After 21 days the mice were sacrificed by cervical dislocation for further analysis. For the cytokine neutralization experiments, A20^myel-KO^ mice were infected with *Trichuris muris* eggs as described earlier and treated with antibodies against IFNγ, IL12 and IL18 (BioXcell, #BE0237) (100μg). The antibodies were administered i.p. every second day for 21 days in total [IFNγ (1mg), IL12 (1mg), IL18 (100μg)]. Antibodies against IFNγ and IL12 were kindly provided by Dr. Louis Boon from Bioceros B.V., Netherlands

### Infection with Salmonella Typhimurium

For infection we used *Salmonella Typhimurium*, virulent strain SL1344. Culture: *Salmonella* from glycerol stock was grown in LB for 1h at 37°C. Later, the liquid culture was plated and grown overnight (ON) on Blood agar plates at 37°C to acquire single colonies. The day before infection (D-1), a) a colony of Salmonella from the plate was inoculated in 5 ml LB and grown ON at 37°C, in a shaking incubator at 180 rpm and b) Treated the mice with Streptomycin to improve colonization efficiency (25mg/mouse). The day of infection (D0), food and water were removed from the cage for 4h. The ON liquid culture was diluted 100x and 500x in LB and grown for 3h at 37°C. OD600 was measure to dermine concentration and make the appropriate dilutions so that mice were infected with 10^7^ CFU Salmonella/200ul. Water and food was returned to the mice and the animals monitored for 8 days for body weight change, temperature andcondition. At the end, the mice were sacrificed and organs processed for further analysis. To evaluate bacterial recovery in spleen, colon and liver we used supplemented MacConkey plates [MacConkey agar (CM0007B; Oxoid), 6.8% Sodium thiosulfate (Sigma), 0.8% Ferric Ammonium citrate (Sigma)].

### Flow cytometry

Isolation of colonic lamina propria of WT and A20^myel-KO^ ice was performed as described previously^42^. The isolated cells were used for extracellular and intracellular staining for representative markers of T and myeloid cells, in two separated panels (1 × 10^6^ cells per sample), then analyzed by flow cytometry using a five-laser – BD LSR Fortessa. For the analysis of the T cell compartment, the cells were stained with 7-AAD (1/1,000) for live/dead separation and then extracellularly with the following antibodies: anti-CD3 APC (eBioscience; 17-0031-83; 1/100), anti-CD4 APC-Cy7 (BD; 552051; 1/200) and anti-CD8 V500 (BD; 560776; 1/100). The cells were then fixed/permeabilized using the Foxp3 Transcription Factor Staining Buffer Set (eBioscience; 00-5523-00). For the intracellular staining the following antibodies were used: anti-Foxp3 Alexa Fluor 488 (eBioscience; 53-5773-82; 1/100), anti-T-bet PE-Cy7 (eBioscience; 25-5825-82; 1/100), anti-RORγt BV421 (BD; 562894; 1/100) and anti-GATA3 PE (eBioscience; 12-9966-42; 1/100). For the analysis of the myeloid compartment, the cells were stained with Fixable Viability Dye eFluor 506 (eBioscience; 65-0866-14; 1/300) for live/dead separation and then only extracellularly with the following antibodies: lineage (CD19 (eBioscience; 15-0193-82; 1/400), CD3 (eBioscience; 15-0031-82; 1/200) and NK1.1 (BioLegend; 108716;1/200) PE-Cy5), anti-CD45 Alexa Fluor 700 (eBioscience; 56-0451-82; 1/800), anti-Ly6G PercpCy5.5 (BD; 560602; 1/200), anti-Ly6C-APC (eBioscience; 17-5932-80; 1/200), or anti-GR-1 where mentioned (eBioscience; 45-5931-80; 1/200), anti-Siglec F BUV395 (BD; 740280; 1/200), anti-CD11b BV605 (BD; 563015; 1/600), anti-CD64 BV711 (BioLegend; 139311; 1/100), anti-F4/80 Biotin (eBioscience; 13-4801-82; 1/100), Streptavidin BV421 (BioLegend; 405226; 1/1000), anti-CD11c PE-eFluor 610 (eBioscience; 61-0114-82; 1/300) and anti-MHC Class II APC-eFluor 780 (eBioscience; 47-5321-80; 1/800). The flow cytometry data were analyzed using FlowJo software and a sequential gating strategy. The graphs showing the percentages of myeloid cells represent proportions of the parent population. t-SNE plots for T cells: 10000 CD3+ cells /genotype were exported and in total 20000 cells analysed. t-SNE plots for myeloid cells: 20000 CD45+ cells (lineage excluded: CD19+, Nk1.1+ and CD3+ fraction)/genotype were exported and in total 40000 cells were analysed. The samples were then concatenated and analyzed with the t-SNE plugin of Flowjo (10.6.1) (iterations = 1,000; perplexity = 30). Following dimensional reduction, coordinates for each t-SNE dimension (i.e., t-SNE1 and t-SNE2) in the two-dimensional plots were determined and integrated as novel parameters. The gating strategies for myeloid and T cell analysis are including at Sup. Fig. 2G and 3.

For the sorting of CD11b+ cells from WT and A20myel-KO mice, the cells were stained with anti-CD11b BV605 (BD; 563015; 1/600) and CD19 (eBioscience; 15-0193-82; 1/400), CD3 (eBioscience; 15-0031-82; 1/200) and NK1.1 (BioLegend; 108716;1/200) PE-Cy5) as lineage markers (sup. Fig. 7A,B).

### Isolation and culture of mesenteric lymph nodes (mLN)

The mesenteric lymph nodes of WT and A20^myel-KO^ mice were harvested postmortem. The tissue was dissociated with a plunger through a 70μm filter. mLN cells were resuspended in RPMI 1640 medium supplemented with fetal calf serum and penicillin/streptomycin and treated with 3μg/ml a-CD3 and 5μg/ml a-CD28, both soluble for 48h. After culture the cells were processed for RNA isolation, cDNA synthesis and quantitative PCR and the supernatants were used for cytokine measurement.

### Ex vivo differentiation of BMDMs

Bone marrow was isolated from tibia and fibula bones of WT and A20myel-KO mice. The samples were treated with ACK for 3 minutes at room temperature to remove red blood cells. The cells were then cultured petri dishes in 10ml RPMI 1640 medium containing 10% fetal calf serum, penicillin/streptomycin and 40ng/ml m-CSF for a total of 7 days. On day three (D3) the cells were supplemented with 1 ml of 400 ng/ml m-CSF. On D5 the medium was replaced with fresh containing 40ng/ml m-CSF. On D7 the differentiated cells were replated and cultured with 4h *Trichuris muris* antigen for different time points i.e. 30min, 1h, 2h, 4h, 6h, 24h, 48h, 72h. At the end of the treatment, the cells were processed for RNA isolation, cDNA synthesis and quantitative PCR and the supernatants were used for cytokine measurement.

### Quantitative polymerase chain reaction (qPCR)

Tissues and cells were lysed using RLT and β-mercaptoethanol and the TissueLyser II (Qiagen, for tissues). Total RNA was isolated using the RNeasy Mini Kit (Qiagen), according to the manufacturer’s instructions. The synthesis of cDNA was performed using QuantiTect® Reverse Transcription Kit (Qiagen). For qPCR, SensiFAST SYBR NO-ROX (BioLine) and specific primers for: GATA3 (fw GGCAGAAAGCAAAATGTTTGCT; Rev TGAGTCTGAATGGCTTATTCACAAAT), IL4 (fw TGGACTCATTCATGGTGCAG; Rev AACATGGGAAAACTCCATGC), IFNγ (Fw TCTGGAGGAACTGGCAAAAG; Rev GCTGATGGCCTGATTGTCTT), IL12 (Fw CCTGGGTGAGCCGACAGAAGC; Rev CCACTCCTGGAACCTAAGCAC) were used on LightCycler 480 (Roche). The reactions were performed in triplicates and the results were analyzed with qbase+ software. As housekeeping genes, GAPDH, Actb and Ubc were used.

### ELISA for IgG1 and IgG2c levels in blood serum

The plates for the ELISA were coated overnight at 4°C with dilute *Trichuris* antigen in 0.05M carbonate bicarbonate buffer. After washing 5x with PBST 0.05%, the plate was incubated with blocking buffer (3% BSA+PBST 0.05%). Diluted serum was added then into the plate and incubated for 90min at room temperature (RT). Later, the plate was washed 5x with PBST 0.05% and incubated with biotinylated antibody G1 (Serotec MCA336B) or G2A (BD bioscience 553388) for 60min at RT. After 5 washes with PBST 0.05%, the plate was incubated with Streptavidin-POD conjugate (Roche, 11089153001) for 1h at RT. Following, the peroxidase substrate ABTS (2,2’-Azinobis [3-ethylbenzothiazoline-6-sulfonic acid]-diammonium salt, Sigma, A1888) was added and the absorbance determined at 405nm, with reference of 490nm.

### Quantification of cytokines

Cytokine levels were determined in mouse blood serum and in supernatants of *ex vivo* cultured mesenteric lymph nodes and bone marrow-derived macrophages. Mouse blood was collected postmortem by cardiac puncture and stored in clot activator tubes containing serum separating silica particles.

IL4 (171G5005M), IL5 (171G5006M) and IL13 (171G5012M) were quantified using the Bio-Plex Pro kit (Bio-Rad) on the Bio-Plex 200 system (Bio-Rad) according to the manufacurer’s instructions. IFNg, IL12 and IL18 were quantified using ELISA (R&D Systems).

### Histology

Mouse colon and caecum tissues were fixed in formalin (10%; neutral buffered; Sigma–Aldrich). Samples were dehydrated, embedded in paraffin, sectioned at 5 μm, stained with hematoxylin and eosin and examined by light microscopy. For combined Alcian Blue and periodic acid Schiff stainings, dewaxed sections were dehydrated and incubated in Alcian Blue for 20 min. The sections were subsequently washed with PBS and treated with 1% periodic acid for 15 min followed by incubation in Schiff’s reagent for 15 min. Sections were counterstained with Mayer’s hematoxylin for 30s, washed and dehydrated before mounting with Depex.

### MACSima platform

The following samples were preparing for MACSima imaging: Colon - WT steady state, Colon – A20 steady state, Colon – WT Trich. Muris, Colon – A20 Trich. Muris. The tissues were mount in OCT compound and sections of 8um thickness were prepared with Cryostat. The sections were later fixed with Acetone, at -20°C for 10min. Washed 3x with PBS and proceed with staining. 32 markers were in total assessed with MACSima technology which were coupled to PE or FITC fluophores. According to the basic principle of the technology, validated antibodies by Miltenyi enable the identification of multiple markers at the same time. The MACSima imaging system acquires fluorescence images of 2 markers/cycle of the desired predefined ROIs. To complete the cycle, the fluorescence signal is erased anf process restarts automatically. The images werer later analysed with MACSiQView software.

### Bulk RNA sequencing and analysis

Sorted CD11b+ cells from colon lamina propria and (ankle and knee) joint synovium were analyzed with bulk RNA sequencing using a Quantseq mRNA library prep on 37 samples and sequencing on the Nextseq500, SR75 by NXTGNT (Faculty of Pharmaceutical Sciences, Ghent University). The quality of the raw data was checked using FastQC (version 0.11.9) and MultiQC (version 1.12). The trimmed data (Trimmomatic version 0.39) was mapped to the mouse reference genome (Assembly: GRCm39, GCA_000001635.9) using STAR (version 2.7.10a). BAM files where created using Samtools (version 1.9) and counted with featureCounts from the Subread package (version 2.0.3).

The quality of the experimental design was assessed using PCA and sample-to-sample distance to determine outliers samples. DESeq2 (version 1.38.2) was used to perform differential expression analysis. P-values were adjusted for multiple testing by DESeq2 using Benjamini-Hochberg correction. Genes were labeled as differentially expressed when the adjusted-p value < 0.05. Functional enrichement was performed using the fgsea (v1.24.0) package to find enriched pathways. A preranked genelist was generated by ordering the shrunk differential expression results according to log2FC. Enrichment results were filtered using a normalized enrichment score (NES) cut-off > 1.

Gene scoring was performed by comparing the expression data of our dataset with published expression data for in vitro M1-classically activate (IFNγ+LPS) macrophages and M2-alternatively activated (IL4) macrophages described by Orecchioni M. et al^18^ This gene list consists of 710 genes identified to be upregulated in classically activate macrophages and 806 genes to be upregulated in alternatively activated macrophages.

### Statistical analysis

Data were analyzed using GraphPad Prism 9.4.1 software. The results are expressed as means ± s.e.m. We used the unpaired Student’s t test for two parameters comparisons or One-way Anova for multiple comparisons. Differences with P < 0.05 were considered statistically significant and denoted as *P < 0.05, **P < 0.01, ***P < 0.001.

## Supporting information

Petta et al. supplementary data

## Acknowledgements

We would like to acknowledge the VIB Animal and Flow Core facility in Ghent for providing expertise and assistance throughout the project. I. Petta is supported by the Belgian Foundation Against Cancer (Stichting tegen kanker) Foundamental postdoc mandate (2021-030). R. Grencis is supported by the Wellcome Trust (Z10661/Z/18/Z). Research in the G. van Loo lab is supported by VIB and research grants from Ghent University (BOF/24J/2021/052 and BOF23/GOA/001) the FWO (G090322N, G026520N, G012618N, EOS-G0H2522N-40007505), the Queen Elisabeth Medical Foundation, the Charcot Foundation, the Belgian Foundation against Cancer (FAF-F/2018/1200), and the FOREUM Foundation for Research in Rheumatology. The Vereecke lab is supported by Ghent University (BOF.GOA031-22, BOF.IBF037-20), FWO (EOS-G0H2522N-40007505) and Foundation Against Cancer (Stichting tegen kanker - F/2020/1421).

