## Supplementary material for "Myeloid A20 is critical for type-2 immune mediated helminth resistance": Petta et al. supplementary data

**Supplementary Figure 1**

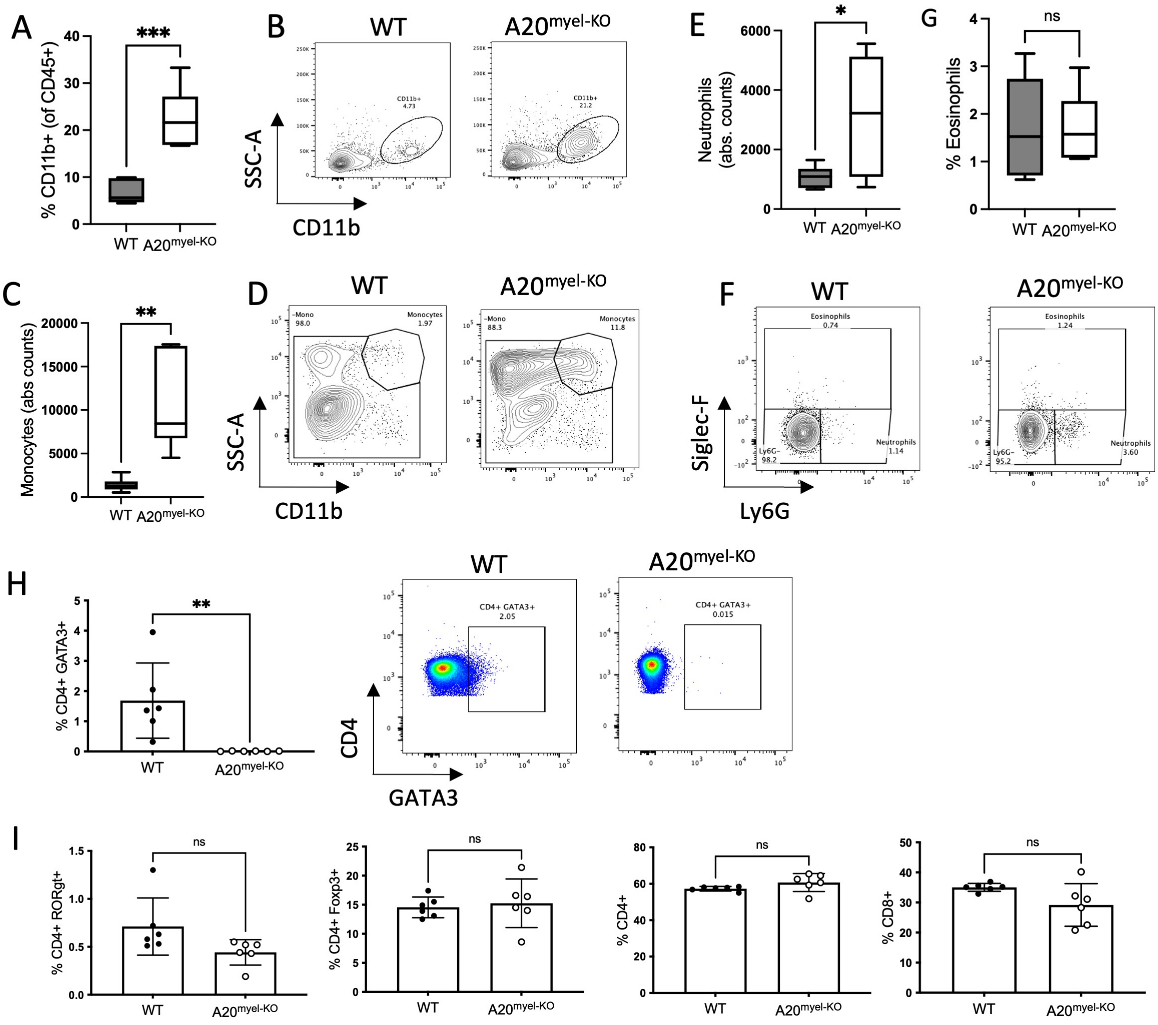


**Supplementary figure 1:** Flow cytometry analysis of spleen of WT (n=6) and A20$^{myel-KO}$ (n=6) mice for the myeloid compartment at steady state, including CD11b+ (A,B), Monocytes (C,D), Neutrophils and Eosinophils (E,F,G). Flow cytometry for T cell analysis in the spleen of WT (n=6) and A20$^{myel-KO}$ (n=6) including (H) CD4+GATA3+, (I) CD4+RORγt+, CD4+Foxp3+, CD4+ and CD8+. Data are presented as means ± s.e.m. Statistical significance was determined by a two-sided Student's t-test.

**Supplementary Figure 2**

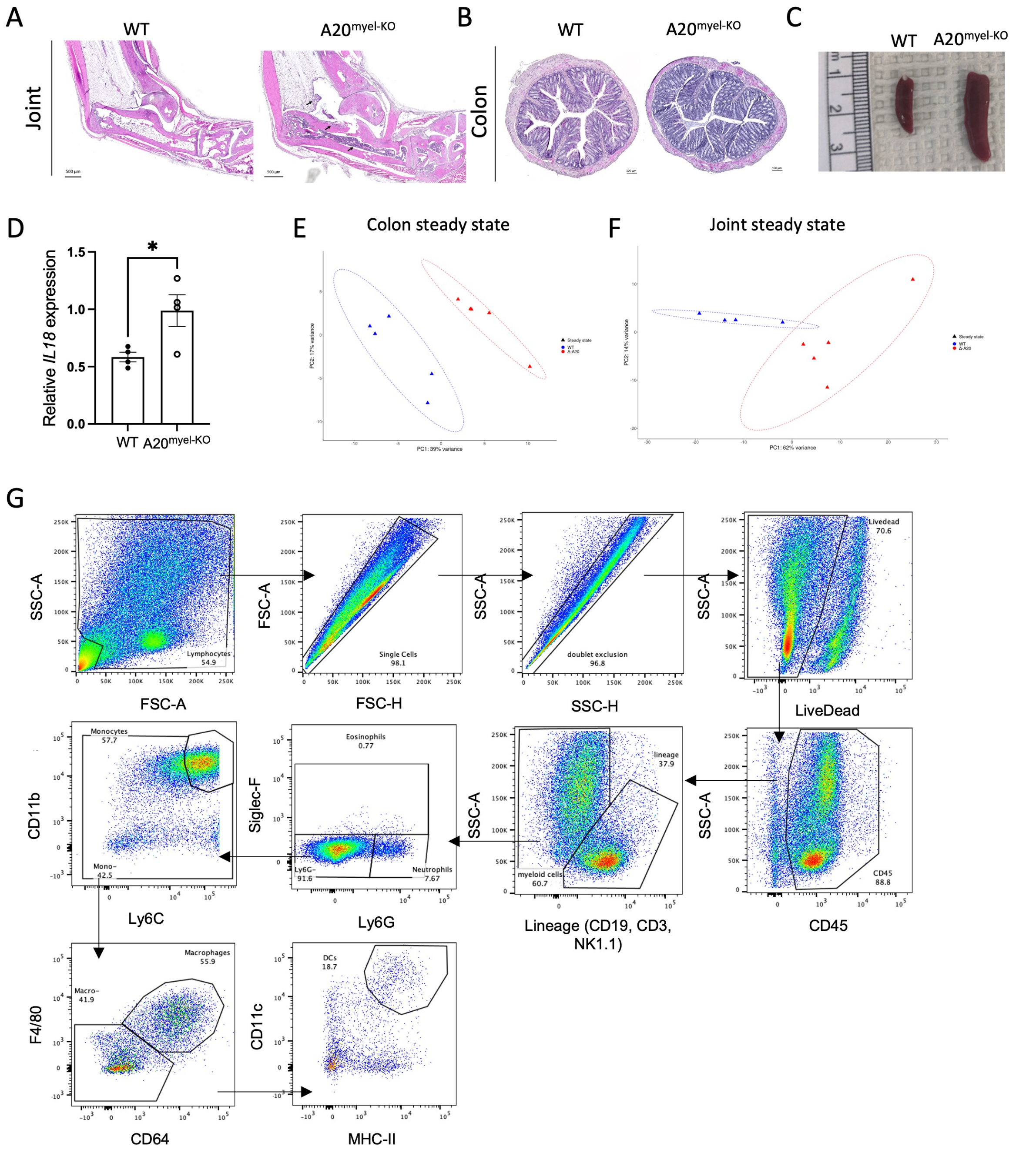


**Supplementary Figure 2:** A) Representative images of H&E staining of ankle joint of WT and A20$^{myel\text{-}KO}$ mice, 14 weeks-old. Arrows indicate the site of immune cell infiltration. B) Representative images of AB/PAS staining of colon sections of WT and A20$^{myel\text{-}KO}$ mice, 14 weeks-old C) Representative macroscopic images of WT and A20$^{myel\text{-}KO}$ spleens after dissection at 14 weeks-old. D) Relative IL18 expression in BMDMs of WT (n=4) and A20$^{myel\text{-}KO}$ (n=4) in steady state condition. Data are presented as means ± s.e.m. Statistical significance was determined by two-sided Student's t-test. E) Bulk RNA sequencing and clustering of WT (n=5)

and A20$^{myel-KO}$ (n=5) colon and F) of WT (n=4) and A20$^{myel-KO}$ (n=5) joint samples on PCA plots according to gene expression. G) Gating strategy for flow cytometry analysis of myeloid cells in colon and spleen.

**Supplementary Figure 3**

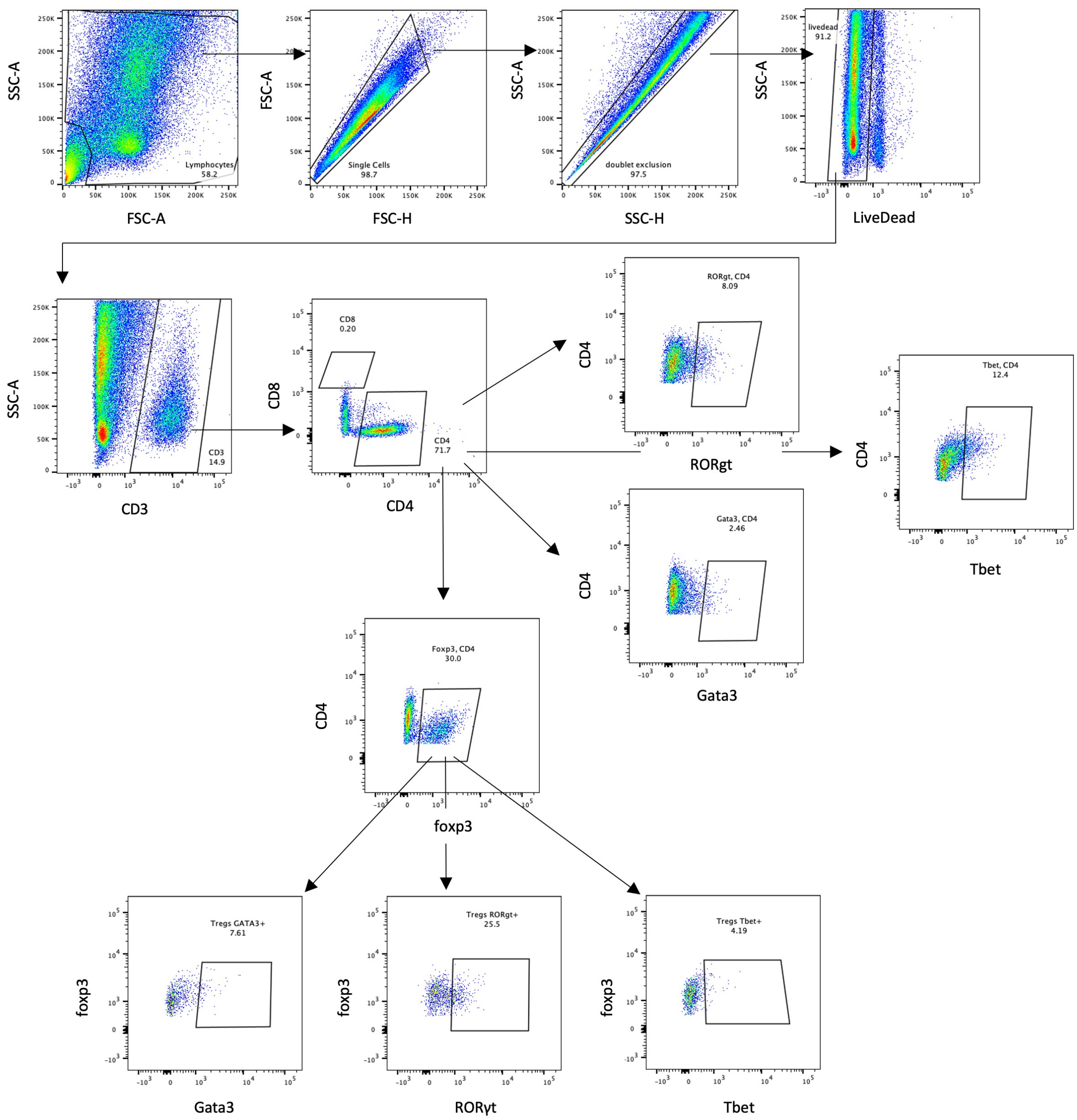


**Supplementary Figure 3:** Gating strategy for flow cytometry analysis data for T cells in colon and spleen. Details for the immune markers analyzed and fluophores used are included in materials and methods section.

**Supplementary Figure 4**

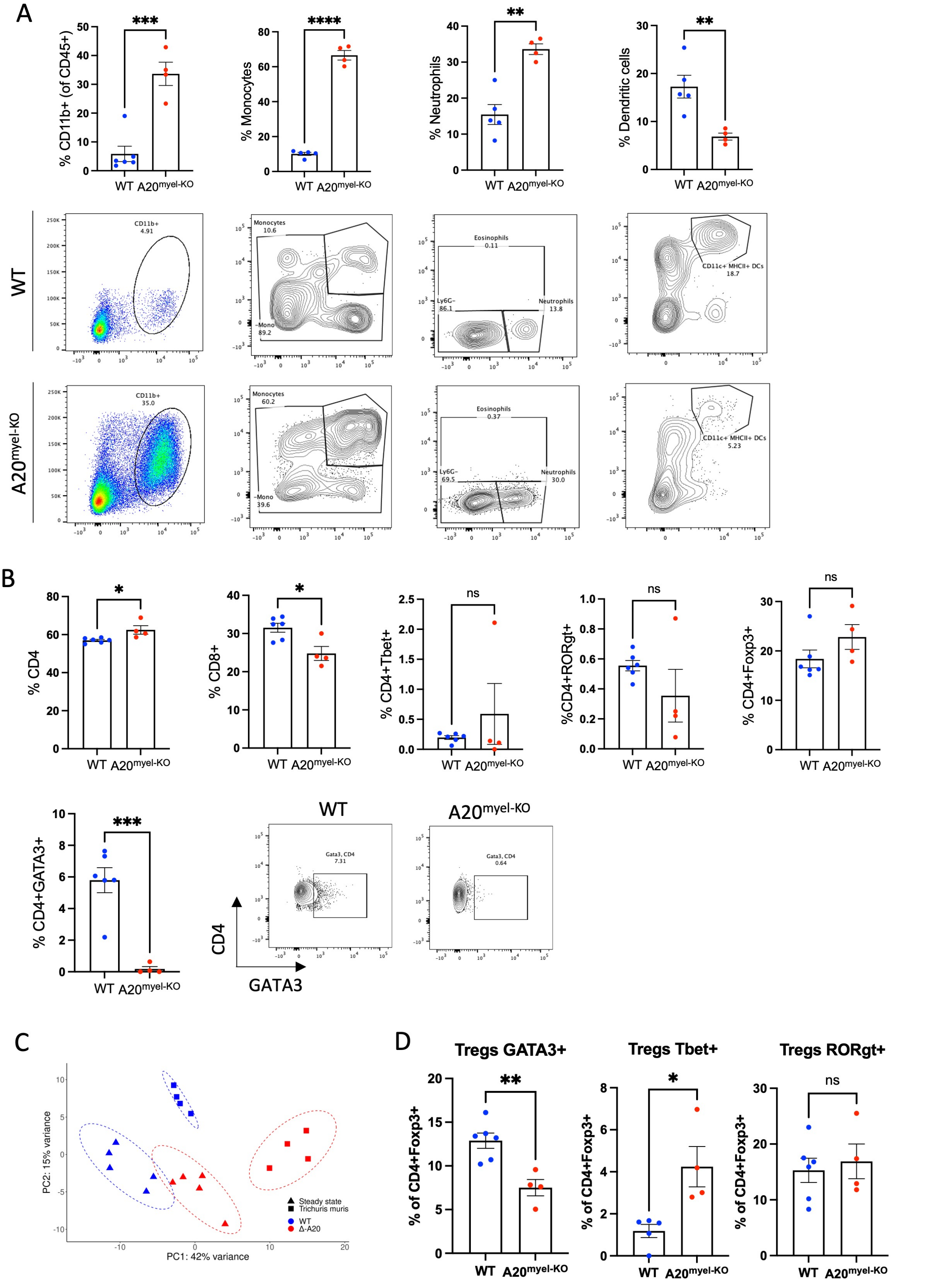


**Supplementary Figure 4:** Flow cytometry analysis in the spleen of WT (n=6) and A20$^{myel\text{-}KO}$ (n=4) upon *Trichuris muris* infection including A) total CD11b+, Monocytes, Neutrophils and

Dendritic cells for the Myeloid compartment. B) Analysis of the T cell compartment includes CD4+, CD8+, CD4+Tbet+, CD4+RORgt+, CD4+Foxp3+ and CD4+GATA3+ levels. C) Bulk RNA sequencing and clustering of WT colon at steady state (n=5, blue triangle) and upon *Trichuris muris* (n=4, blue square) versus A20$^{myel-KO}$ colon at steady state (n=5, red triangles) and upon *Trichuris muris* infection (n=4, red squares) combined on a PCA plot according to gene expression. D) Tbet, GATA3 and RORγt expression within colonic Treg population in WT (n=6) and A20$^{myel-KO}$ (n=4) upon Trichuris muris infection.

**Supplementary Figure 5**

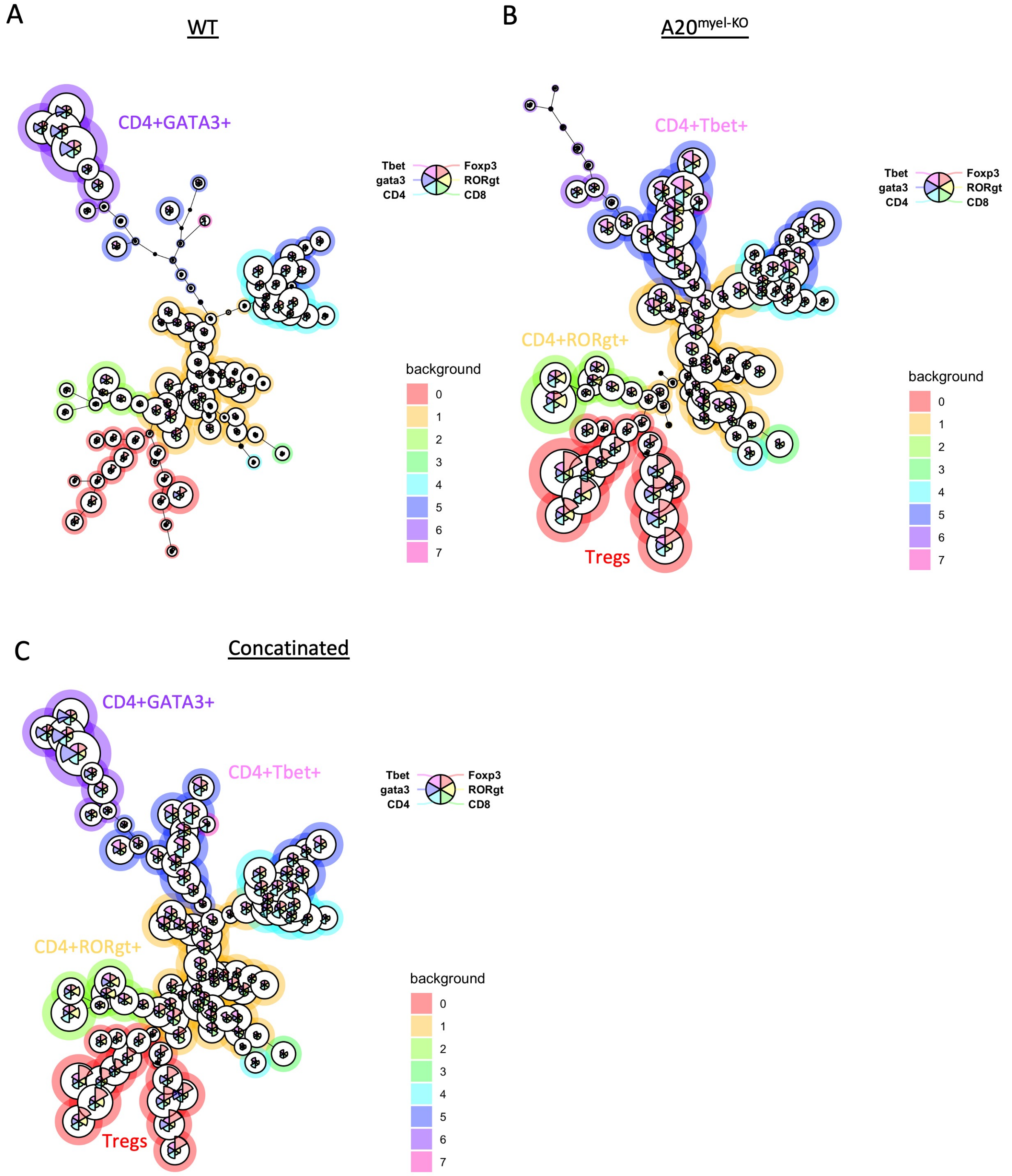


**Supplementary Figure 5:** Flowsom analysis of WT (n=4) and A20[myel-KO] (n=4) upon *Trichuris muris* infection for T cell composition. The data are visualized in minimal spanning trees and according to the parameters expression levels, Flowsom organizes the branches of the tree (cell clusters) in 8 different metaclusters (0-7). The parameters analysed are colored inside every circle as following: Tbet=magenta, GATA3=purple, CD4=light blue, CD8=green,

RORγt=yellow and Foxp3=orange. The height of each part indicates the expression intensity: the higher the part in the circle (node), the higher the expression of the marker is. The size of the circles is also indicative of the abundance of the representative cell cluster. A) In WT mice upon Trichuris infection we observe an abundance of the cell cluster which is represented by circles that contain higher purple parts (GATA3). In contrast, A20$^{myel\text{-}KO}$ mice show abundance of the cell cluster which is represented by circles that contain higher magenta parts (Tbet). We also observe increase of the circles contain higher yellow parts (RORγt). Tregs are represented by orange parts and appear as a heterogenous population separated in 2 branches, with higher expression of RORγt (yellow), Tbet (magenta) and GATA3 (purple). The samples were exported and concatenated and a total of 10.000 CD3+ cells/genotype is analysed. The parameters applied for the Flowsom analysis are the following: Number of metaclusters=8, SOM (self-organising map) grid size 10x10, Node scale=300%. Flowjo plugin 3.0.18[43]. The different metaclusters present the differential and combinatorial expression of the analysed parameters. 5200 CD3+ cells were extracted and concatenated/genotype (in total 10400 cells) to perform Flowsom analysis.

**Supplementary Figure 6**

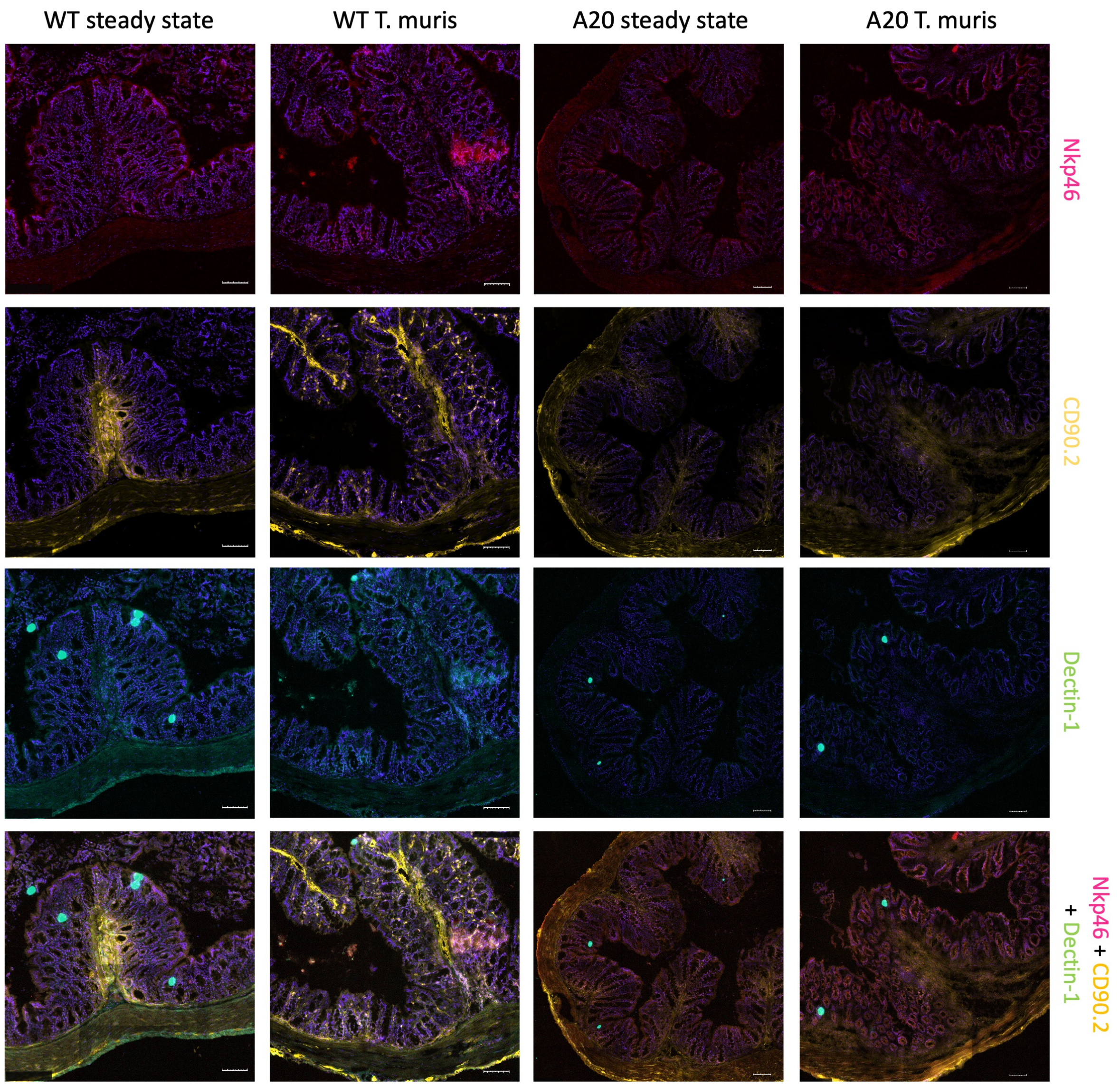


**Supplementary Figure 6:** MACSima fluorescent imaging of colon WT and A20$^{myel\text{-}KO}$ in steady state and upon *Trichuris muris* infection for Innate Lymphoid cells (ILCs). ILC1 (Nkp46, magenta), total ILCs (CD90.2, yellow), ILC3 (Dectin-1, blue). Scale bars 100µm.

**Supplementary Figure 7**

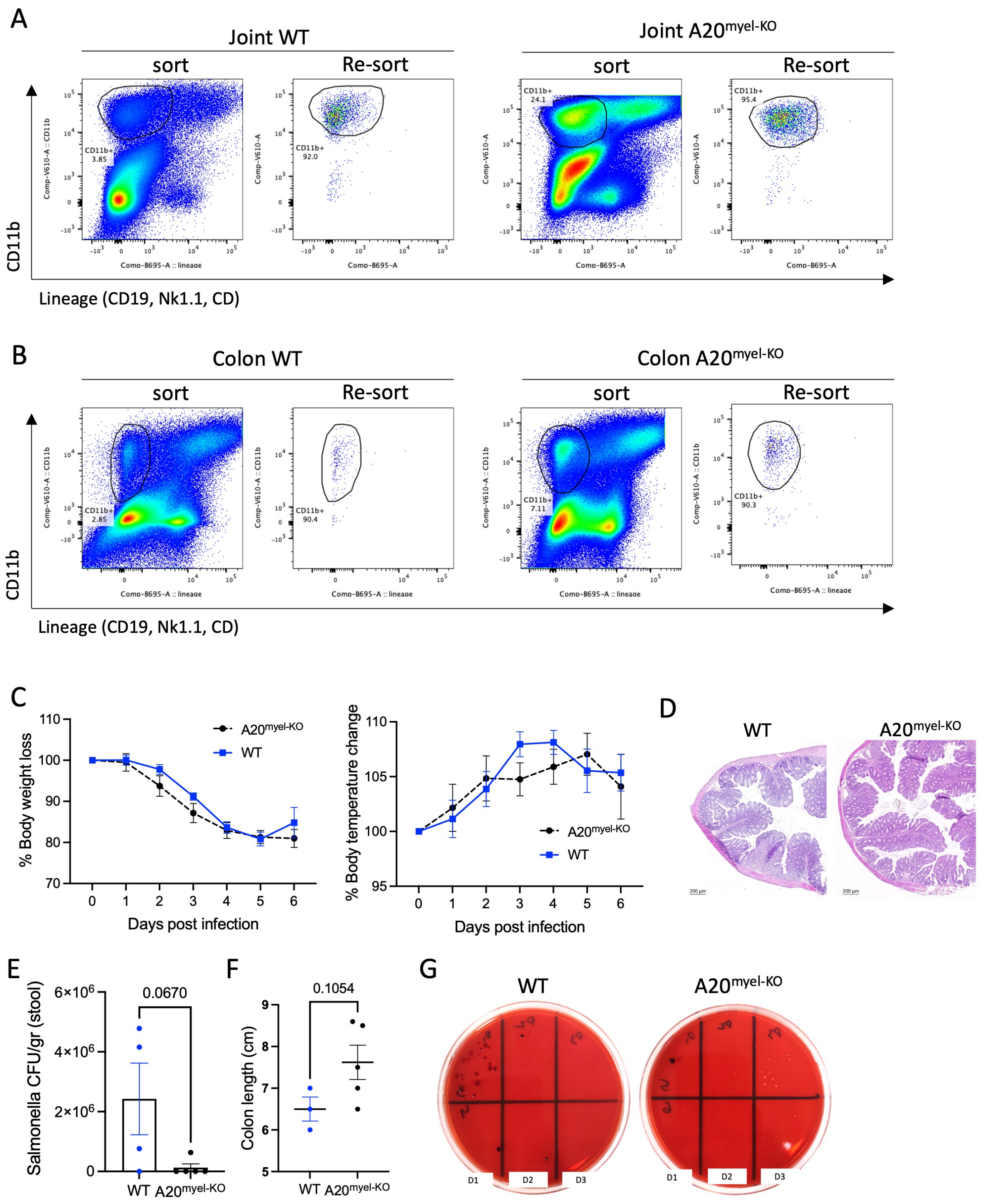


**Supplementary Figure 7:** Fluorescence-activated cell sorting of CD11b+ from (A) joint synovium and (B) colon lamina propria of WT and A20$^{myel-KO}$ mice for bluk RNA sequencing. Representive plots of sorted and re-sorted DC11b+ cells to ensure the purity of the sorted population. C) Graphs presenting the % body weight loss (left) and % body temperature change (right) in WT (blue) and A20$^{myel-KO}$ (black dotted) mice upon infection with Salmonella

Typhimurium. D) Representative H&E staining of colon section of WT and A20$^{myel\text{-}KO}$ on day 8 p.i. E) CFU counts of retrieved *Salmonella Typhimurium* from stool samples of infected WT (n=4) and A20$^{myel\text{-}KO}$ (n=5). F) Colon length of infected WT (n=3) and A20$^{myel\text{-}KO}$ (n=5) 8 days post infection. G) McConkey plate showing *Salmonella Typhimurium* colony growth in WT (n=2/plate) and A20$^{myel\text{-}KO}$ (n=2/plate) from liver samples on day 8 p.i. The samples are serially diluted (1/10) left to right (D1->D3).

### Supplementary Figure 8

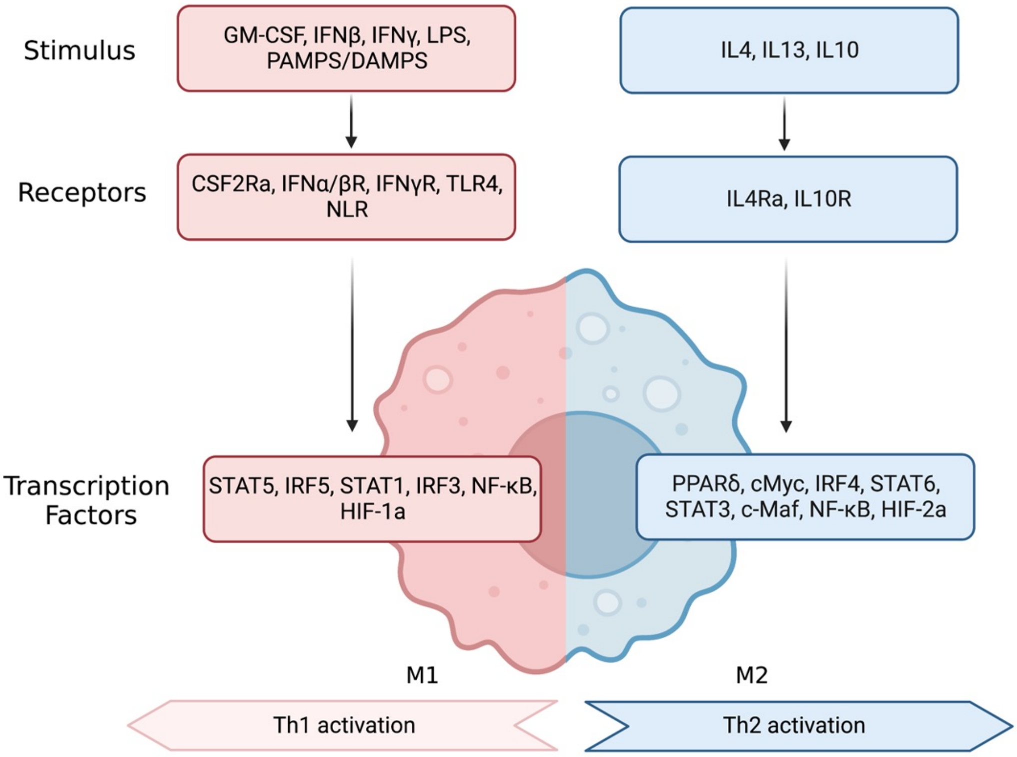


**Supplementary Figure 8:** Scheme of Macrophage polarization towards M1 or M2 phenotype including stimuli, receptors and transcription factors involved.
